# Neither alpha-synuclein-preformed fibrils derived from patients with *GBA1* mutations nor the host murine genotype significantly influence seeding efficacy in the mouse olfactory bulb

**DOI:** 10.1101/2023.08.24.554646

**Authors:** Sara Walton, Alexis Fenyi, Tyler Tittle, Ellen Sidransky, Gian Pal, Solji Choi, Ronald Melki, Bryan A. Killinger, Jeffrey H. Kordower

## Abstract

Parkinson’s disease (PD) is a neurodegenerative disease characterized by progressive motor symptoms and alpha-synuclein (αsyn) aggregation in the nervous system. For unclear reasons, PD patients with certain GBA mutations (GBA-PD) have a more aggressive clinical progression. Two testable hypotheses that can potentially account for this phenomenon are that *GBA1* mutations promote αsyn spread or drive the generation of highly pathogenic αsyn polymorphs (i.e., strains). We tested these hypotheses by treating homozygous *GBA1* D409V knockin (KI) mice with human α-syn-preformed fibrils (PFFs) and treating wild-type mice (WT) with several αsyn-PFF polymorphs amplified from brain autopsy samples collected from patients with idiopathic PD and GBA-PD patients with either homozygous or heterozygous *GBA1* mutations. Robust phosphorylated-αsyn (PSER129) positive pathology was observed at the injection site (i.e., the olfactory bulb granular layer) and throughout the brain six months following PFF injection. The PFF seeding efficiency and degree of spread were similar regardless of the mouse genotype or PFF polymorphs. We found that PFFs amplified from the human brain, regardless of patient genotype, were generally more effective seeders than wholly synthetic PFFs (i.e., non-amplified); however, PFF concentration differed between these two studies, and this might also account for the observed differences. To investigate whether the molecular composition of pathology differed between different seeding conditions, we permed Biotinylation by Antibody Recognition on PSER129 (BAR-PSER129). We found that for BAR-PSER129, the endogenous PSER129 pool dominated identified interactions, and thus, very few potential interactions were explicitly identified for seeded pathology. However, we found Dctn2 interaction was shared across all PFF conditions, and Nckap1 and Ap3b2 were unique to PFFs amplified from GBA-PD brains of heterozygous mutation carriers. In conclusion, both the genotype and αsyn strain had little effect on overall seeding efficacy and global PSER129-interactions.

## Introduction

Parkinson’s disease (PD), the second-most common neurodegenerative disorder after Alzheimer’s Disease, affects nearly one million people in the U.S., with the number expected to rise to 14 million worldwide by 2040^1^. PD is characterized by motor symptoms, including bradykinesia, postural instability, tremor, rigidity^2–4^, and non-motor symptoms, including constipation, urinary dysfunction, depression, anosmia, psychosis, apathy, and sleep disorders. The development of these symptoms is associated with the accumulation of misfolded alpha-synuclein (αsyn) protein into intracellular inclusions called Lewy pathology (LP)^5–7^.

Several known genetic risk factors exist for PD, with mutations in *GBA1* being the most common. *GBA1* encodes the lysosomal enzyme glucocerebrosidase (Gcase), which is deficient in Gaucher disease. The first indication of a link between parkinsonism and *GBA1* mutations stems from observations of PD in patients with Gaucher disease and in their relatives that were carriers^8–11^. Patients with *GBA1*-associated PD (GBA-PD) tend to have an earlier age of onset and, with certain mutations, exhibit faster motor and cognitive decline than patients without the mutation^12^. These patients also show rapid accumulation and spread of αsyn pathology^13^. Several hypotheses have been proposed to explain patients with GBA-PD have such rapid clinical decline and widespread αsyn accumulation. *GBA1* loss-of-function (LOF) coupled with the pathological spread of αsyn results in decreased lysosomal activity and function, altered αsyn processing, and accumulation of pathologic αsyn, could contribute to the observed aggressive disease^14–17^. Thus, aggressive αsyn spread and clinical progression could result from reduced Gcase enzymatic activity. Alternatively, the αsyn strain or polymorph associated with *GBA1* mutations could be particularly virulent, as observed with αsyn strains purified in the presence of detergents from MSA brain^18^.

Several *GBA1* mutant mice have been developed, including *GBA1* D409V knock-in mice which have the human *GBA1* D409V point mutation inserted into the mouse *Gba1* gene. *GBA1* D409V mice have reduced GCase activity and accumulation of glycosphingolipid in the brain and liver^19^. Although de novo asyn pathology is not observed in the brain of these mice^19^ they are more susceptible to asyn accumulation^20^, have increased asyn expression at 12 months of age^20,21^, and show may enhance spread via increased exocytosis of asyn^21^.

Here we investigate whether the aggressive progression of clinical symptoms in GBA-PD patients results from loss of GCase activity or the unique polymorphs formed in the GBA-PD brain.

## Results

We bilaterally injected *de novo* assembled αsyn-PFFs into the granular layer (GL) of the olfactory bulb (OB) of *GBA1* D409V KI mice and separately injected PFFs seeded by pathologic αsyn isolated from brain homogenates from patients with GBA and idiopathic (i.e., without known genetic cause) PD (GBA-PFFs and idiopathic PD-PFFs, respectively) into the OB of WT mice (See Fig. 1 for summary). For GBA-PFFs, fibrils from heterozygous *GBA1* (*GBA1*^+/-^) carriers with PD were amplified from the cingulate cortex (cases #GBA1, GBA3, GBA7). The homozygous *GBA1* (*GBA1*^-/-^) variant PFFs were amplified from the frontal cortex (cases #14-388 and 12-373), and idiopathic variant (*GBA1*^+/+^) PFFs were amplified from the cingulate cortex (B19-Cingulate). Before fibril amplification, the amount of pathogenic aggregated and phosphorylated αsyn was measured in brain homogenates using a filter retardation assay and Fluorescence resonance energy transfer (FRET) assay (Fig. S1). Seeding reaction kinetics were monitored with thioflavin T (Fig. S2), and fibril morphology was assessed by TEM and limited proteinase K (PK) digestion (Fig. S3). The resulting PFFs were injected bilaterally into the OB GL based on previously published protocols^22^. All samples were pretreated with PK to enhance αsyn pathology detection and avoid detecting endogenous PSER129 (Fig. 2A). PK treatment dramatically reduced endogenous PSER129 staining and revealed otherwise undetected αsyn pathology (Fig. 2A, “GL”)

**Figure 1.**
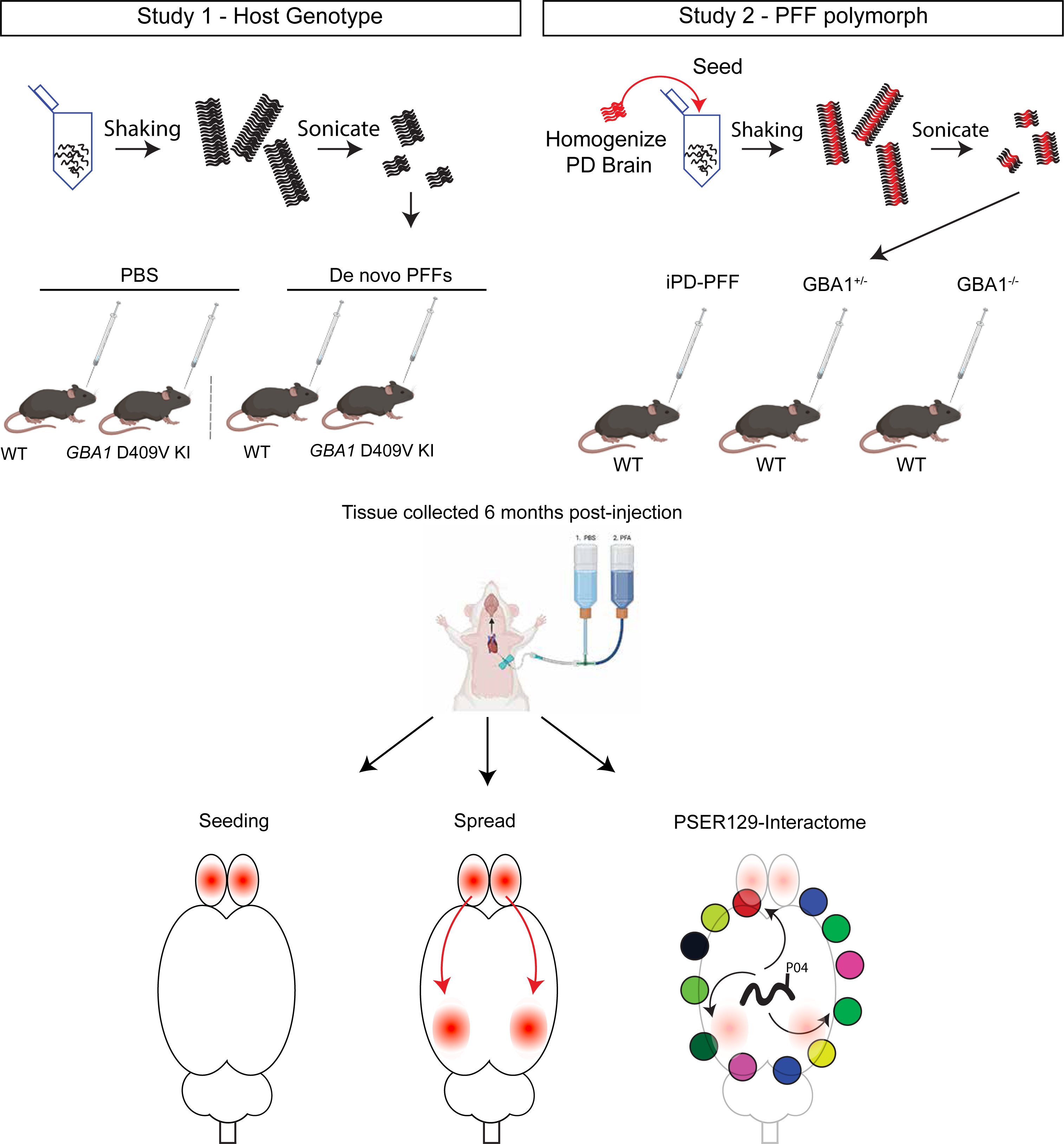
Summary of approach. Two studies were conducted. Study 1, GBA1 D409V KI mice or WT mice were bilaterally injected with de novo αsyn PFFs or PBS into the olfactory bulb (OB) granular layer. Study 2, PFFs were first generated in the presence of brain homogenates from idiopathic PD patients (iPD), or PD patients with heterozygous or homozygous *GBA1* mutations. The resulting PFF polymorphs were then injected bilaterally into the OB of WT mice. Mice were sacrificed at 6 months post OB-injection and perfused with PBS followed by 4% PFA. Seeding and spread of αsyn pathology was determined with IHC detection of PSER129. Global PSER129 interactions were determined using BAR-PSER129.

**Figure 2.**
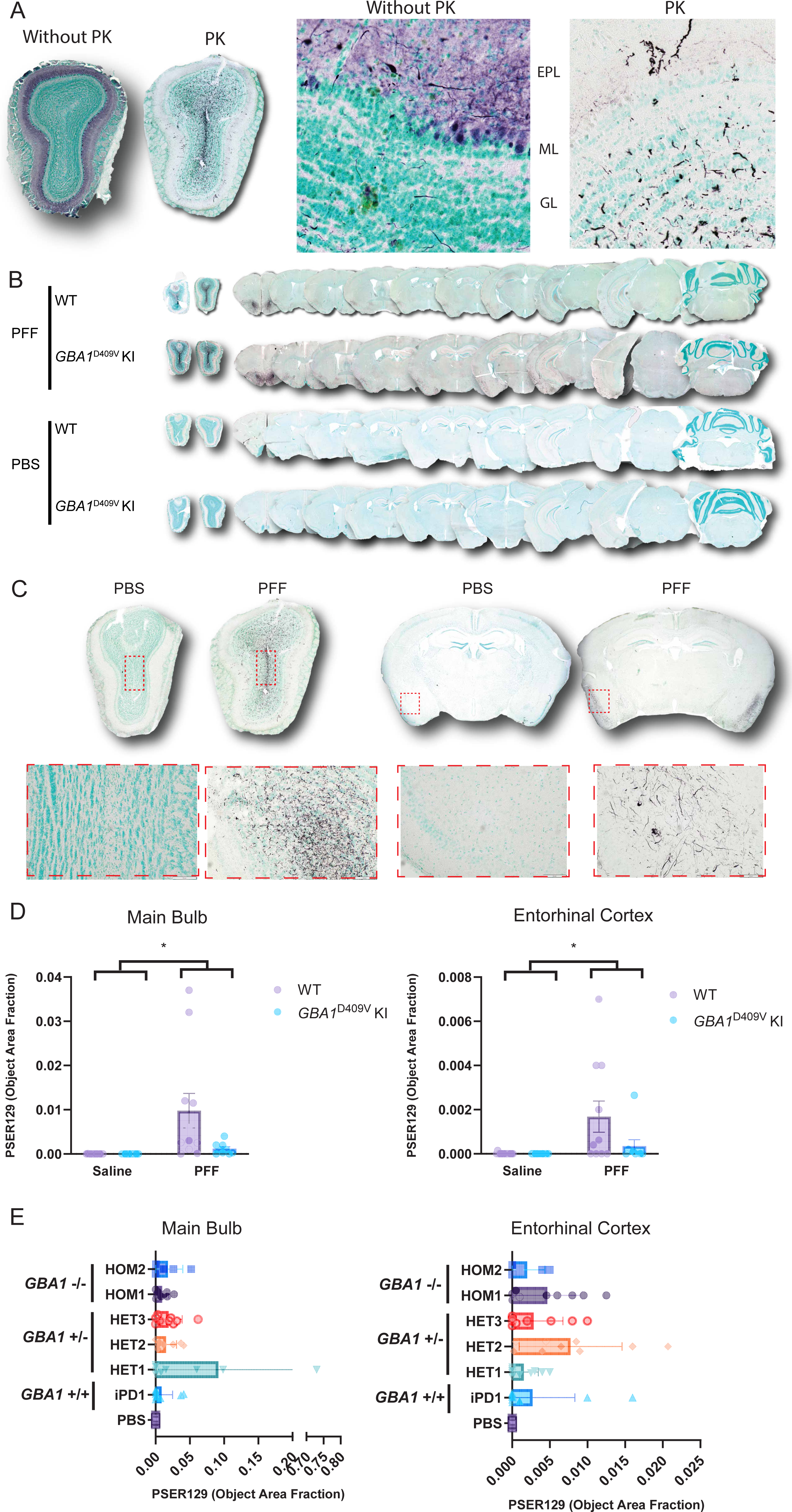
Seeding and spreading following PFF injection into mouse OB. (A) Differentiation of PSER129-positive pathology from endogenous PSER129 pool required pretreatment of tissues with proteinase K (PK). OB sections from a single PFF-injected mouse were stained for PSER129, with or without PK pretreatment. High magnification images show PSER129 reactivity in the granular layer (GL), mitral cell layer (MT), and external plexiform layer (EPL). (B) Pathological asyn distribution across the neuroaxis 6 months following bilateral injections of PFFs into 4-month-old WT and *GBA1*D409V KI mice. Tissues were treated with PK prior to immunostaining PSER129 using TSA. Sections counterstained with methyl green, making nuclei appear light green-blue. PSER129 was detected using the chromogen nickel-enhanced DAB, which appears black. Representative whole-section scans show the tissue distribution of PSER129 positive asyn pathology throughout the brain. (C) Representative images of OB and Entorhinal cortex showing PSER129 staining that was quantified. Red dotted box superimposed on whole tissue scans highlights the approximate areas that the high magnification image was taken. (D) Quantification of PK-PSER129 reactivity in the main OB and entorhinal cortex of WT and *GBA1*D490V KI mice treated with PFFs. (E) Quantification of PK-PSER129 reactivity in the main OB and entorhinal cortex of mice treated with asyn PFF polymorphs amplified from PD and GBA-PD clinical brain specimens. *Two-Way ANOVA F (1, 40) = 7.235, P=0.0104. n = 5-14.

### PSER129 throughout the neuroaxis after PFF seeding

Following bilateral PFFs injection into the OB GL, we observed pathology throughout the neuroaxis (Fig. 2B). We made similar observations as previously reported^22^ and detected PSER129-positive pathology mainly concentrated in several brain regions, including the olfactory bulb (OB), anterior olfactory nucleus (AON), and entorhinal (EC), with little or rare PSER129-positivity in other brain regions. (Fig. 2B, C). We did not observe PSER129 positivity in the SN or STR of any PFF-treated mouse, regardless of host genotype or PFF polymorph. PBS-treated mice showed very little PSER129 staining, with only rare weakly reactive cells (e.g., hippocampus and OB mitral cell layer), which may have resulted from insufficient PK digestion or αsyn ’s interaction with cellular lipid components^23^. Early tests confirmed that endogenous and seeded PSER129 could not be differentiated without PK digestion (Fig. 2A).

We confirmed, as previously reported^22^, that 6 months after OB-PFF injections the majority of αsyn pathology was detected in the main OB and the EC (Fig. 2B, C). Therefore, to determine αsyn pathology spread, we quantified PSER129 in the main OB and EC (Fig. 2C, D). Results showed a significant increase (EC, Two-way ANOVA, F(1,40)=7.235, p=0.0104; OB, Two-way ANOVA, F(1,39)=7.327, p=0.01) in PSER129 in both brain regions when comparing PBS vs. PFFs (Fig. 2D). However, we did not observe any statistically significant differences between WT and *GBA1*^D409V^ KI mice. We observed OB-PFF injections in both *GBA1*^D409V^ KI and WT mice resulted in a variable abundance of PSER129 pathology (Fig 1D, WT, mean =0.00168, SD= 0.00234, n =11; GBA1^D409V^ mean = 0.00035, SD = 0.001, n = 9). Some animals displayed profound pathology and others minimal pathology consisting of a few observable PSER129-positive neurites in both brain regions. Similarly, following treatment with PFF-polymorphs, we observed a significant increase (EC, Two-way ANOVA, F(1,40)=7.235, p=0.0104; OB, Two-way ANOVA, F(1,39)=7.327, p=0.01) in PSER129 in the main OB and EC; interestingly, on average, an order of magnitude greater than non-strain PFFs (Fig. S4). However, the apparent difference in PSER129 pathology abundance between de novo generated and amplified PFFs was non-significant (Students t test, t (70) = 1, p = 0.3207) (Fig S4). PSER129 pathology abundance was not significantly different between distinct PFF strains, in the main OB or in the EC.

### PSER129 in the OB following PFF seeding

PFF injections into the OB GL can propagate via mitral cell axons to olfactory cortical areas or back-propagate through mitral cells dendrites to the outer plexiform and glomerular layers. Several lines of evidence suggest that different PFF-polymorphs or host mutations promote cell specificity for the spreading process, resulting in a different pattern of spread and, thus, different presentation of clinical disease symptoms^24^.To determine if either genotype or PFF strains influenced the direction of spread from a common injection site in the OB GL, we measured PSER129 abundance across the layers of the OB (Fig. 3A). Results showed that abundant PSER129 was consistently detected in the GL (i.e., injection site) in the OB of PFF-treated mice (Fig 2B, C). The second most abundant site of detection was the mitral cell layer. One PFF-treated *GBA1*^D409V^ KI mouse showed very high PSER129 abundance in the mitral cell layer. We observed the glomerular layer and outer plexiform PSER129 following PFFs, but these two layers typically had very low levels of pathology. PBS-treated mice did not have detectable PSER129 in any OB layer, regardless of genotype (Fig. 3B). PFF-strain-treated mice showed significantly more PSER129 in the GL when compared to other OB layers (Fig. 3C). No significant differences were observed between patient derived PFFs (Fig. 3C).

**Figure 3.**
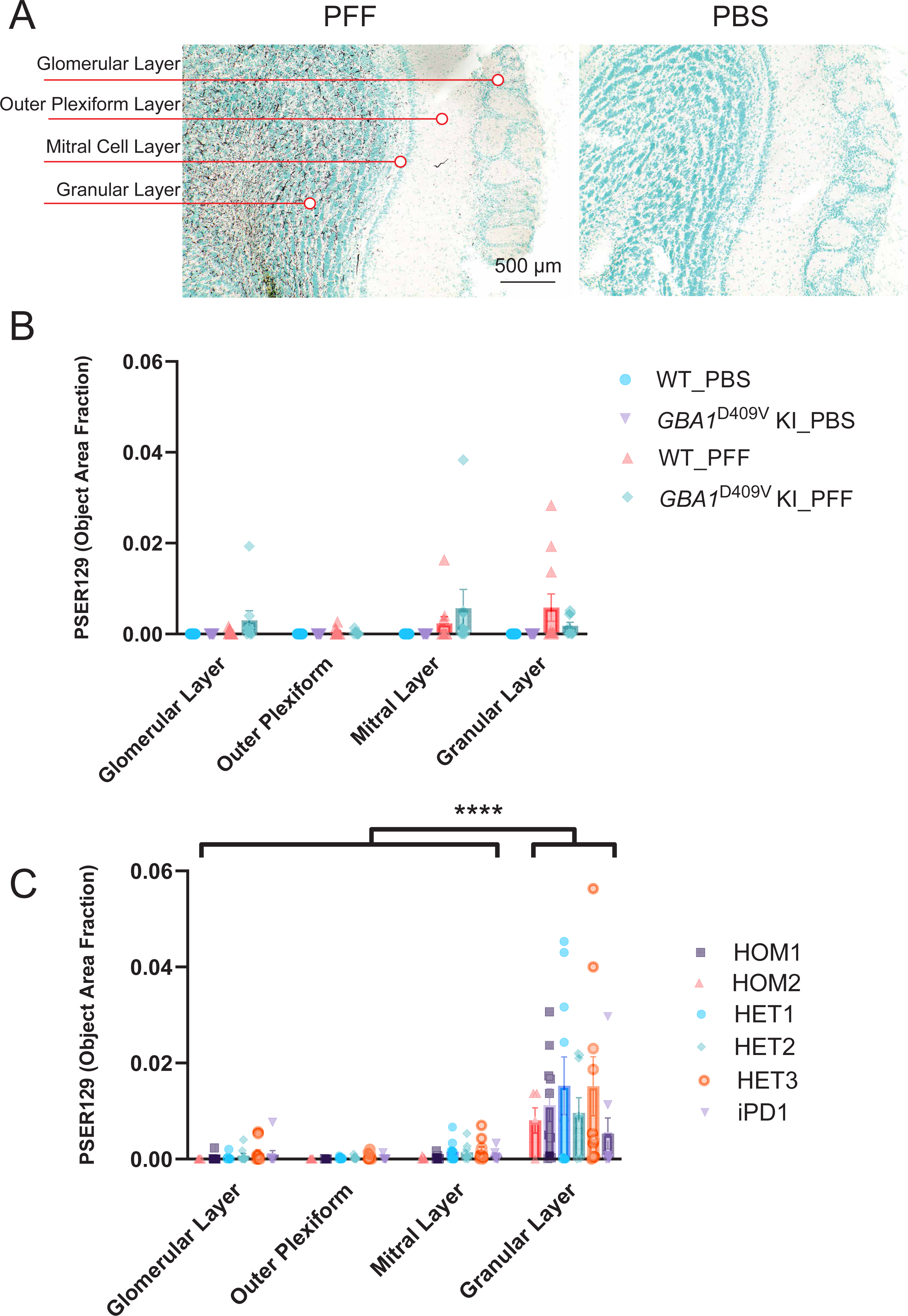
Distribution of asyn pathology across OB layers following PFF injection into mouse OB. (A) Representative images of PFF and PBS injected OB showing distribution of PK-PSER129 reactivity across OB layers. Each layer quantified is annotated to the left of the images. (B) Quantification of PK-PSER129 across the layers of WT and *GBA1*D409V KI mice seeded with PFFs. (C) Quantification of PK-PSER129 across the layers of WT mice treated with various strains of asyn PFFs amplified from PD and GBA-PD clinical specimens. **** F (1.021, 47.97) = 26.42, P<0.0001. n = 5-14.

### BAR

Seeding and spread (distribution) were similar in *GBA1*^D409V^ KI mice and between PFF polymorphs. Next, we investigated whether the molecular makeup of seeded pathology was similar between the different seeding conditions by performing BAR-PSER129 on pooled samples from each experimental group. BAR-PSER129 is a high-throughput method to identify protein-protein interactions^23,25,26^. Results showed diffuse BAR-PSER129 labeling throughout many brain regions, including cortical areas and the hippocampus (Fig. 4A), consistent with the endogenous PSER129 pool^23^. Mice exposed to PFFs showed dense BAR-PSER129 labeling primarily concentrated near the site of PFF injection (i.e., OB GL) and the entorhinal cortices. In many brain regions, the endogenous PSER129 signal was very intense and obscured the detection of PFF-seeded pathology that was observed using proteinase K protocols (Fig. 2 and Fig. 3). Western blot (WB) of extracted proteins from all samples showed remarkably similar total αsyn and PSER129 content across all samples tested regardless of PFF exposure (Fig. 4B). This suggests that despite observing PK-resistant PSER129 positive αsyn pathology in the brain, overall, αsyn pathology only makes up a small proportion of the total brain αsyn pool (∼5% or less). Following BAR-PSER129 capture enriched proteins were spotted onto a PVDF membrane and probed for several markers (Fig. 4C). Biotin, PSER129, and αsyn content were all enriched when comparing BAR-PSER129 and BAR-NEG (antibody omission control) capture conditions (Fig. 4C). BAR-PSER129 enrichment was observed for pooled samples from “polymorphs” and “genotype” study groups (Fig. 4D and 3E, respectively). BAR-PSER129 showed higher enrichment for PSER129 in animals seeded with PFF polymorphs than all other treatment groups (34-fold vs. 2.9-fold, respectively). Together, results show that BAR-PSER129 labeling enriched for both endogenous PSER129 and PSER129 positive inclusions in mouse brains.

**Figure 4.**
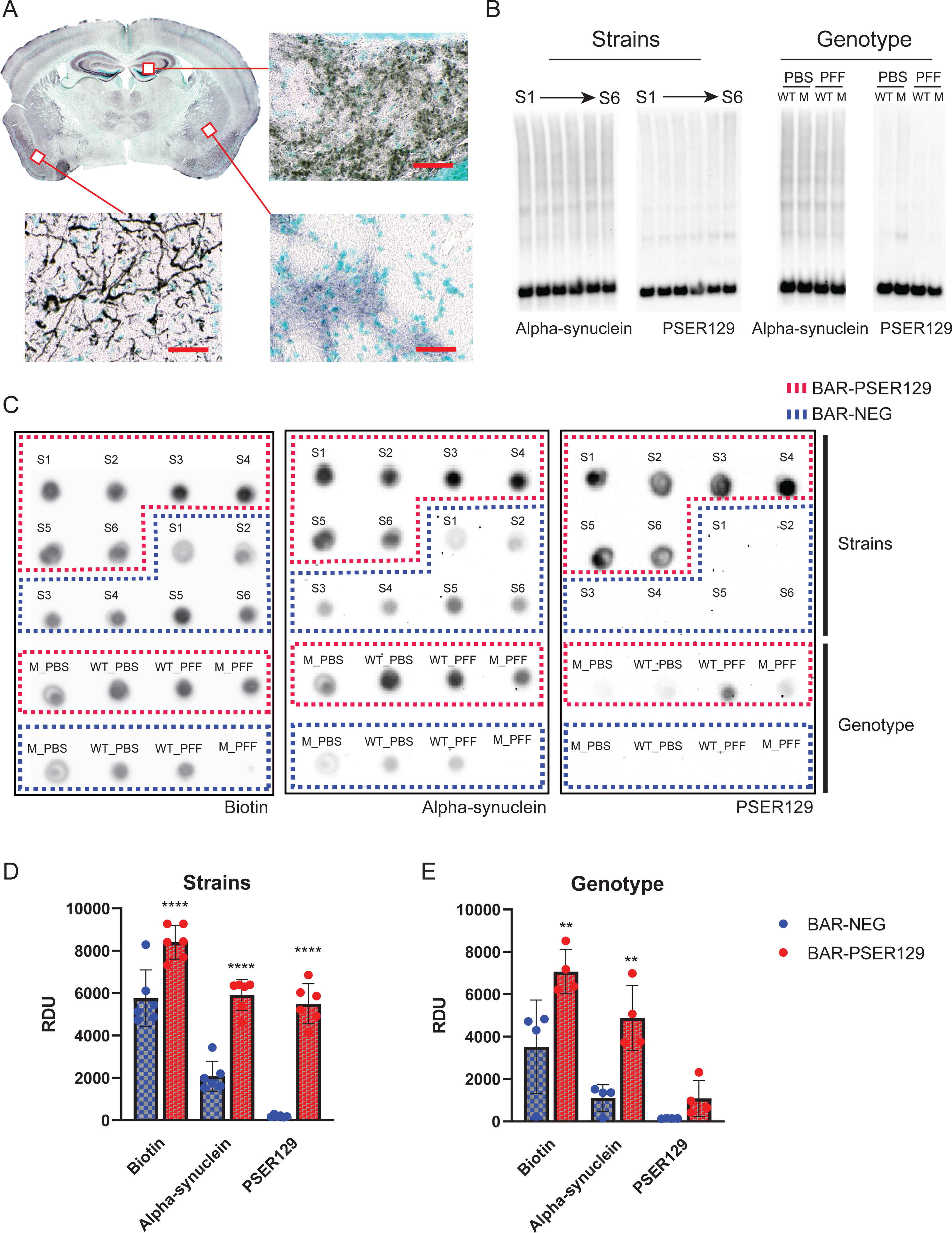
BAR labeling and enrichment for PSER129 interacting proteins for mice under different seeding conditions. BAR-PSER129 was performed on pooled brain sections from mice treated with varying strains of PFFs and PFFs seeded into mice carrying the *GBA1*D409V mutation. (A) Representative BAR-PSER129 labeled brain section. BAR labeled proteins are visualized using ABC reagent and nickel DAB. Enlarged images show BAR-PSER129 labeled both an endogenous PSER129 signal and seeded pathology. Bottom panel shows PSER129 pathology in the Piriform areas, while other two panels show endogenous PSER129 signal in the amygdala and hippocampus. (B) Western blot of proteins extracted from non-BAR labeled brain sections. Blots labeled “Strains” are blots of proteins extracted from animals seeded with six (S1-S6) different PFF strains. “Genotype” are blots of proteins extracted from WT and *GBA1*D409V KI mutant mice (“M”) treated with PBS or PFFs. Both total asyn and PSER129 were detected. (C) Dot blot of BAR-PSER129 and BAR-NEG (i.e., without antibody, background) captured proteins. Blots were probed for biotin, asyn, and PSER129. The position of capture conditions are denoted on the blots. (D) (E) Densitometry analysis of dot blots showing enrichment for biotin, asyn, and PSER129. Graphs show the enrichment for samples from Strain and Genotype studies. All blots are presented uncut. Each tissue pool = 4-5 mice. **** p<0.0001 Two-way ANOVA, ** p<0.001 Two-way ANOVA, Šídák’s multiple comparisons test.

Next, we performed liquid chromatography-tandem mass spectrometry (LC-MS/MS) on BAR fractions and identified a total of 464 proteins across all samples. Unbiased hierarchical clustering revealed samples were segregated primarily by capture conditions, BAR-PSER129 or BAR-NEG (Fig. 5A). Paired multiple t-test revealed that 136 proteins were enriched (FDR < 0.01) over background (i.e., BAR-NEG) including Snca, Sncb, Atp6v1e1, Syn1 (FDR < 3 X 10^-6^) (Fig. 5B, Fig. S6 Summary of BAR-PSER129 identified proteins). SNCA was the most abundantly enriched protein along with several presynaptic/SNARE proteins (Fig. S5). STRING pathway analysis of all statistically significant BAR-PSER129 enriched proteins highlights known interactions between BAR-PSER129-identified proteins (Fig. 5C). Snca was a central node of the interaction map occurring within the major interaction cluster (MCL clustering, inflation parameter = 1.6) that was significantly enriched for thousands of pathways including “SNARE complex disassembly”, “Synuclein”, and “Parkinson’s disease.” Other functional clusters distinct from Snca were enriched for “Glycolysis”, “Synaptic vesicle endocytosis”, “Parkinson Disease/Ubiquitinating process”, “Vacuolar proton-transporting v-type atpase complex”, and “mRNA splicesome.” The PSER129 interaction map identified here is largely consistent with previously reported αsyn interactomes and αsyn hypothesized SNARE functions^27,28^. We determined all proteins significantly enriched with BAR-PSER129 regardless of treatment condition. Next, we evaluated whether any PSER129-interactions were exclusive or specific to seeding conditions (i.e., αsyn strain or *GBA1* mutations). Of all 464 proteins, 9 were found to be exclusively detected in mice treated with PFFs (Fig. 5D) and not in PBS treated mice, WT or *GBA1*^D409V^ KI. STRING analysis showed that these proteins were functionally distinct as only a single functional interaction was found between Ran and Cd3. Dctn2 was found in nearly all PFF seeding conditions with the exception of PFF seeded *GBA1*^D409V^ KI mice. No enriched pathways were found for this set of identified proteins. None of the identified PFF specific proteins were found to be significantly enriched with BAR-PSER129, overall.

**Figure 5.**
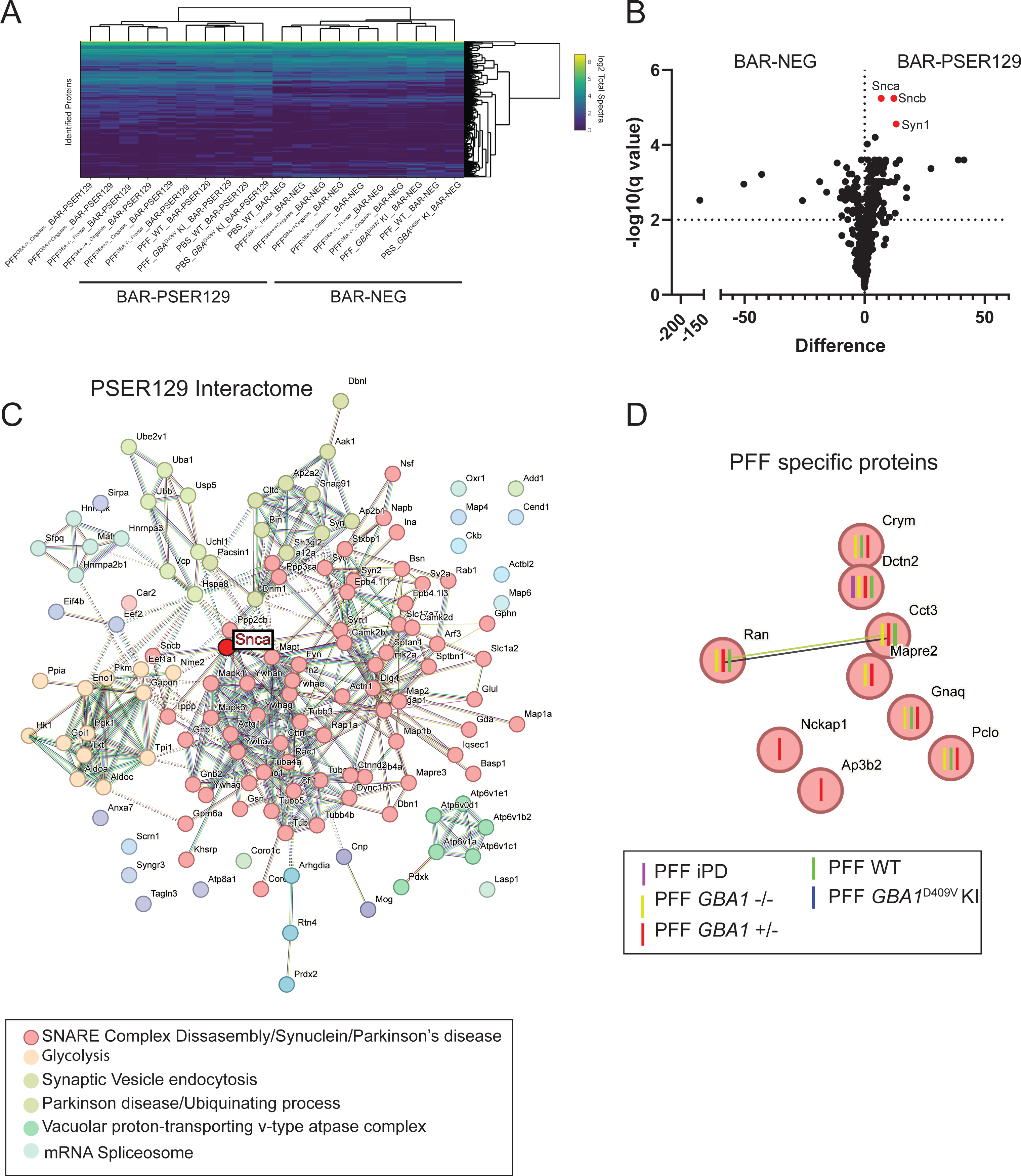
Identified PSER129 interactome in the brain of mice under different seeding conditions. BAR-PSER129 was performed on pooled (4-5 mice) brain samples from each experimental group. LC-MS/MS identified a total of 464 proteins. (A) Unbiased hierarchical clustering of identified proteins (A). Two major clusters were observed according to capture conditions and annotated “BAR-PSER129” and “BAR-NEG.” (B) Volcano plot of all proteins identified. BAR-PSER129 and BAR-NEG were compared. Top hits, Alpha-(Snca), beta-synuclein (Sncb), and synaptophysin I (Syn1) were denoted in red and labeled. (C) STRING interaction map of 144 proteins significantly enriched over background (BAR-NEG). MCL clustering (inflation parameter 1.4) grouped proteins into 6 functional clusters. Most populated clusters are annotated with consensus terms for significantly enriched pathways. The position of asyn (Snca) is highlighted. (D) PSER129-interacting proteins identified exclusively in brains of PFF seeded mice. Each protein is annotated with the seeding conditions for which the protein was identified. “PD PFF”, “GBA+/+ PFF”, “GBA-/- PFF” are PFF strains derived from PD cases without GBA mutations, homozygous, and heterozygous mutations, respectively. “PFF WT” and “PFF GBA1D409V” were WT and *GBA1* D409V KImice, respectively, seeded with PFFs.

## Discussion

Several recent reports have investigated whether *GBA1* mutations influence αsyn PFF seeding behavior and have reached conflicting conclusions^20,29–31^. Here we took the unique approach of assessing both host-genotype and unique αsyn PFF polymorphs, to account for the aggressive clinical outcomes seen in GBA-PD. Following OB-PFF injections, αsyn pathology was predominantly observed in the GL and the EC. However, we did not observe any significant differences for seeding and spread between αsyn polymorphs or in GBA1^D409V^ KI mice seeded with de novo αsyn PFFs. Although we assessed these animals’ behavior, we did not observe evidence of impaired motor skills or impaired olfaction across treatment conditions (Fig. S7-14). These findings broadly agree with previous observations that PFF and OB-PFF seeding models typically do not result in overt disease-relevant behavioral phenotypes^22,32^. Furthermore, non-progressive spread^22^ or the lack of involvement of crucial upstream/downstream disease pathways might account for the lack of phenotypes observed in PFF models. Our findings support the conclusion that in αsyn PFF models, neither strains from patients with *GBA1* mutations, nor host *murine Gba1 genotype* influence seeding efficiency or spread^20,30^. Instead, *GBA1* mutations might act upstream on the endogenous αsyn pool creating a cellular environment conducive to αsyn misfolding and aggregation^20,31,33^. αsyn PFFs are useful for modeling downstream processes (i.e., seeding/spread) but not upstream processes (i.e., *de novo* αsyn misfolding/aggregation) and this might account for the lack of influence of *GBA1* mutations in αsyn PFF models. For example, *GBA1* mutations might promote αsyn pathology generation by creating a lipid environment detrimental to αsyn vesicle interactions resulting in the probability of αsyn misfolding and accumulation^34–36^. Alternatively, LOF *GBA1* mutations can increase αsyn expression in neurons^21^, which would increase the probability of de novo αsyn aggregation or seeding/spread^31^.

OB-PFFs result in widespread PK resistant PSER129 pathology throughout the neuroaxis. Our data is largely consistent with previous descriptions of the OB-PFF model^22,32^. However, a notable exception, we found that striatum and substantia nigra did not develop PSER129-pathology up to 6 months following PFF injection. Instead, we observed robust pathology in brain nuclei directly innervated with mitral cell axons, but pathology was limited to secondary or tertiary brain regions. This supports the conclusion that αsyn pathology spreads via mitral cell axons but progressive spread (i.e., secondary, and tertiary cells) is minimal.

The IHC protocol used here improved αsyn pathology detection in our PFF model. Optimized PK-digestion removed endogenous PSER129 while tyramide signal amplification allowed for high sensitivity detection of PK resistant-PSER129 (Fig. 2A). The resulting high signal-to-noise ratio allowed us to unambiguously detect and differentiate αsyn pathology from the endogenous PSER129 pool, something that had not been done prior. An interesting resulting observation was the apparent lack of pathology in OB mitral cells and their apical dendrites, which normally contain a high concentration of endogenous PSER129, and are the high probability conduit for PFF spread from the OB. The reason (s) for this observation are not entirely clear.

There are several limitations for these studies. First, PK-resistant PSER129 was used to measure αsyn pathology and other pathology markers (e.g., PK-resistant αsyn, conformation specific antibodies) were not explored. Second, BAR-PSER129 identified PSER129-interactome was dominated by endogenous PSER129, and αsyn pathology was only a minor component. Although this endogenous PSER129 pool agrees with recent reports^37,38^ it made identification of unambiguous PFF-specific or genotype specific PSER129-interactions via BAR difficult. Future studies should.

## Conclusion

In conclusion, we did not observe marked differences between the seeding and spread of αsyn in *GBA1*^D409V^ KI mice or for αsyn PFFs amplified from GBA or idiopathic PD brain homogenates. Overall PSER129-interactome was unaffected by PFF seeding, however, a few identified PSER129-interacting proteins were exclusive to PFF seeded brains and specific PFF polymorphs. Neither the *GBA1*^D409V^ mutation or αsyn PFF strain affect seeding efficacy and αsyn pathology spread, but rather they likely affect upstream αsyn turnover, leading to its accumulation and subsequent aggregation.

## Methods

### PFF generation and characterization

Human full-length WT αsyn (canonical sequence) was expressed in *E. coli* BL21 DE3 CodonPlus cells and purified as previously described^39^. The monomeric state of the protein was assessed by analytical ultracentrifugation^40^ and the absence of contaminating endotoxins was checked using the Pierce LAL Chromogenic Endotoxin Quantification Kit ^41^. *De novo* assembled PFFs were generated as described^40^. PFFs amplified from patient brain homogenates were obtained as previously described^42^ for detailed case information see Table S1. All assembly reactions were monitored by thioflavin T binding and the nature of the fibrillar assemblies was assessed by transmission electron microscopy after negative staining with 1% uranyl acetate and limited proteolysis using proteinase K followed by PAGE (Fig. S3).

All PFFs were centrifuged twice at 15,000 g for 10 min and re-suspended twice in PBS. Their concentration was adjusted to 350µM in PBS. They were then fragmented to an average length of 40–50 nm by sonication for 20 min in 2 mL Eppendorf tubes using a Vial Tweeter powered by an ultrasonic processor UIS250 v (250 W, 2.4 kHz; Hielscher Ultrasonic), 6µl aliquots were flash frozen in liquid nitrogen and stored at - 80°C^43^. Before injections, the PFFs were thawed and gently dispersed using a cup horn sonicator (QSonica; 15% power, for 10 pulses of 0.5 s ON, 1 s OFF, repeated four times with 5-min intervals).

### Stereotactic injections

All surgical procedures were done as previously described^22^. *GBA1^D^*^409^*^V^*^/D409V^ mice and WT mice (3 months old) were anesthetized by intraperitoneal injection of ketamine/xylazine mixture (100mg/kg and 10mg/kg, respectively) and stereotactically injected with sterile PBS, PFFs, GBA-PPFs or PD-PFFs into the OB. The stereotactic injections were preformed using a 10 µl Hamilton microsyringe with a 26-gauge needle. For each injection, 1 µl of PFF solution (5 µg/µl for PFFs and 1 µg/µl for αsyn polymorphs) or PBS was bilaterally injected into the OB (coordinates: AP, +5.4 mm; ML, +/- 0.75 mm; DV, −1 mm relative to bregma and dural surface) at a constant rate of 0.2 µl per minute. The needle was left in place for 5 min after injection, and then slowly removed.

### Mouse tissue preparation

Mice were sacrificed 6 months following PFF injection. Mice were anesthetized with ketamine/xylazine (100mg/kg and 10mg/kg, respectively) and transcardially perfused with 0.9% saline, followed by fixation with 4% paraformaldehyde (PFA) in phosphate buffer. Brains were collected, post-fixed for 24 h in 4% PFA at 4°C, and equilibrated in successive sucrose solutions (i.e., 15% then 30% sucrose in phosphate buffer). Brains were stored at 4°C until they were sectioned. Each mouse brain was sectioned into 40-µm free-floating coronal sections on a freezing microtome and stored in a cryoprotectant solution () at 4°C until downstream processing.

### Animals

180 three-month old homozygous C57BL/6N-Gba1tm1.1Mjff/J mice (*GBA1* D409V KI) and C57BL/6 WT mice were purchased from Jackson Laboratories. Animal maintenance and experiments were performed in accordance with the National Institutes of Health guidelines and approved by the Institutional Animal Care and Use Committee of the Rush University Medical Center (Chicago, IL). Mice were maintained at room temperatures of 65–75 °F (∼18–23 °C) with 40–60% humidity. Mice were kept on a 14/10 h light/dark cycle and given a continuous supply of food and water.

### Immunohistochemistry

Immunohistochemistry was performed essentially as described previously^26^. Fixed floating sections were mounted onto gelatin coated slides and dried at room temperature overnight. Slides were rehydrated in TBST (20mM Tris-HCl pH 7.4, 150 mM NaCl, 0.05% triton X 100) and digested with proteinase K (PK, 20 µg/mL) diluted in TBST for 20 min at 37°C. Slides were then fixed in 4% paraformaldehyde for 20 min, rinsed 3 times in TBST, and incubated with 3% hydrogen peroxide for 30 min to quench endogenous peroxidases. Slides were placed in blocking buffer (TBST, 3% bovine serum albumin, 2% goat serum) for 1h and then incubated overnight at 4°C in blocking buffer containing anti-PSER129 antibody EP1536Y (Abcam) diluted 1:50,000. The next day slides were washed 3 times in TBST and incubated with biotinylated anti-rabbit antibody (Vector Labs) diluted 1:400 in blocking buffer for 1h. Slides were washed 3 times in TBST and incubated with ABC reagent diluted in blocking buffer for 1h. Slides were washed twice with borate buffer (0.1M Sodium tetraborate pH 8.5) and incubate in borate buffer containing 0.003% hydrogen peroxide and 5µM biotinyl tyramide (SigmaAlrich) for 30 min. Slides were washed 3 times in TBST and incubated with ABC reagent for 1h. Slides were then washed in TBST and developed using nickel enhanced DAB as previously described ^44^. Slides were counterstained with methylgreen (Sigma), dehydrated with graded alcohols, cleared with xylenes, and cover slipped with cytoseal 60 (Fisher Scientific). Brightfield microscopy was performed using Nikon A1 laser scanning microscope. Density analysis including binary masks and R01 analyses were performed using Elements software (Nikon).

### BAR sample preparation

BAR was performed essentially as previously described ^23,26,45^. Briefly, brain sections collected at 240-micron intervals across the neuroaxis were placed into a net well (Brain research laboratories) and washed 3 times for 1 hour each in TBST. Sections were then placed in 0.3% hydrogen peroxide and 0.1% sodium azide diluted in blocking buffer for 1h at room temperature to quench endogenous peroxidases. Sections were then briefly rinsed in TBST and incubated in anti-PSER129 antibody EP1536Y diluted 1:50,000 in blocking buffer overnight at 4°C with gentle agitation. The following day, sections were washed 3 times in TBST, then incubated with biotinylated anti-rabbit antibody diluted 1:200 in blocking buffer for 1h at room temperature. Sections were then washed 3 times in TBST, incubated with ABC reagent for 1h, and washed off with borate buffer. Sections were then incubated with borate buffer containing biotinyl tyramide as described above. Sections were then washed overnight with TBST, gathered in a 1.5mL Eppendorf tube, centrifuged at 3,000 X g for 15 min to pellet floating sections, and supernatant discarded. Each sample was then briefly sonicated in 1mL of crosslink reversal buffer (5% SDS, 500mM Tris-HCl pH 8.0, 150mM NaCl, 2mM EDTA) and heated for 30 minutes at 98°C followed by 1h at 90°C. Samples were centrifuged at 20,000 x g for 20 min and the supernatant then diluted 1:10 in modified TBST (20 mM Tris-HCl, 200mM NaCl, 2mM EDTA, and 0.5% Triton X-100). Each sample was then incubated with 40mg of streptavidin magnetic beads (Thermofisher Scientific) for 2h at room temperature with constant mixing. Beads were collected using a magnetic stand (Thermofisher Scientific), beads were washed 3 times in 10 mL modified TBST, and then overnight in 10mL of stringent wash buffer (20mM Tris-HCl pH 7.6, 200 mM NaCl, 0.1% SDS, 2mM EDTA). The following day beads were collected using magnetic stand and resuspended in 100 µl 1 X Bolt LDS sample buffer with reducing agent (Thermofisher) then heated for 10 min at 98°C. Samples were vortexed vigorously and beads removed using magnetic stand. 70µl of the sample was electrophoresed approximately 2 cm into a Bolt gel (ThermoFisher). The gel was then fixed in 50% ethanol and 10% acetic acid for 1h. The gel was washed several times in dH20, and proteins stained with colloidal Coomassie blue. The entire sample was then excised for trypsin digestion and mass spectrometry.

### Mass Spectrometry

Samples were prepared and LC-MS/MS conducted as previously described ^23^. Briefly, gel pieces were washed with 100 mM ammonium bicarbonate (AmB)/acetonitrile (ACN) and reduced with 10 mM dithiothreitol (DTT) at 50°C for 45 minutes. Cysteines were alkylated using 100 mM iodoacetamide in the dark for 45 minutes at room temperature (RT). Gel bands were washed in 100 mM AmB/ACN prior to adding 1 µg trypsin (Promega #V5111) for overnight incubation at 37°C. Peptide containing supernatants were collected into a separate tube. Gel pieces were washed with gentle shaking in 50% ACN/1% FA at RT for ten minutes, and supernatant was collected in the previous tubes. Final peptide extraction step was done with 80% ACN/1% FA, and 100% ACN, and all supernatant was collected. Peptides were dried in a speedvac and reconstituted with 5% ACN/0.1% FA in water before injecting into LC-MS/MS.

Peptides were analyzed by LC-MS/MS using a Dionex UltiMate 3000 Rapid Separation nanoLC coupled to an Orbitrap Elite Mass Spectrometer (Thermo Fisher Scientific Inc.). Samples were loaded onto the trap column, which was 150 μm x 3 cm in-house packed with 3 µm ReproSil-Pur® beads. The analytical column was a 75 µm x 10.5 cm PicoChip column packed with 3 µm ReproSil-Pur® beads (New Objective, Inc. Woburn, MA). The flow rate was kept at 300 nL/min. All fractions were eluted from the analytical column at a flow rate of 300 nL/min using an initial gradient elution of 5% B from 0 to 5 min, transitioned to 40% over 100 min, 60% for 4 mins, ramping up to 90% B for 3 min, holding 90% B for 3 min, followed by re-equilibration of 5% B at 10 min with a total run time of 120 min. Mass spectra (MS) and tandem mass spectra (MS/MS) were recorded in positive-ion and high-sensitivity mode with a resolution of ∼60,000 full-width half-maximum. The 15 most abundant precursor ions in each MS1 scan were selected for fragmentation by collision-induced dissociation (CID) at 35% normalized collision energy in the ion trap. Previously selected ions were dynamically excluded from re-selection for 60 s. The collected raw files spectra were stored in. raw format.

Proteins were identified from the MS raw files using the Mascot search engine (Matrix Science, London, UK. version 2.5.1). MS/MS spectra were searched against the SwissProt mouse database. All searches included carbamidomethyl cysteine as a fixed modification and oxidized methionine, deamidated asparagine and aspartic acid, and acetylated N-terminal as variable modifications. Three missed tryptic cleavages were allowed. A 1% false discovery rate cutoff was applied at the peptide level. Only proteins with a minimum of two peptides above the cutoff were considered for further study. Identified peptides/protein were visualized by Scaffold software (version 5.0, Proteome Software Inc., Portland, OR).

### Spot Blotting

To estimate BAR enrichment prior to LC-MS/MS, 1 µl of bead eluent was applied to a methanol activated polyvinylidene difluoride (PVDF) membrane and then allowed to dry completely. The membrane was then reactivated in methanol, rinsed with water, and post-fixed in 4% PFA for 30 min. Blots were then rinse with TBST (20mM Tris-HCl pH 7.6, 150mM NaCl, 0.1% Tween-20) and blocked with buffer containing either BSA (TBST and 5% BSA) or non-fat milk (TBST and 5% non-fat milk) for detection of biotin or αsyn, respectively. Biotinylated proteins were detected by ABC (VectorLabs) diluted 1:10 in BSA blocking buffer for 1 h at room temperature. Αsyn was detected using SYN1 (BD Biosciences) diluted 1:2,000 and PSER129 detected using EP1536Y diluted 1:50,000 both diluted in non-fat milk blocking buffer. Primary antibodies were detected by incubating blots for 1h in secondary anti-mouse HRP conjugate diluted 1:6,000 or secondary anti-rabbit HRP conjugate (Cell signaling) diluted in milk blocking buffer. Following secondary antibody, membranes were washed in high stringency wash buffer (20mM Tris-HCl pH 7.6, 400mM NaCl, 0.1% Tween-20) and imaged using enhanced chemiluminescence (ECL) substrate (Biorad, product # 1705060) and Chemidoc imager (Biorad).

### Western Blotting

Proteins were extracted from fixed floating tissue sections as described above. Proteins were precipitated via chloroform-methanol (61) and dissolved 5% SDS. Protein concentrations was determined using bicinchoninic acid assay (BCA, ThermoFisher). 20 micrograms of total protein was then separated on 4-12% BOLT gel (ThermoFisher). Proteins were blotted onto activated PVDF membranes, transferred proteins were then fixed by placing the membranes in 4% PFA for 30 min. Membranes were then rinsed in water, dried completely, and reactivated in methanol for immunoblotting. Membranes incubated 1 h in blocking buffer (TBST with 5% non-fat milk, 0.5% polyvinylpyrrolidone) and then with either SYN1 (diluted 1:2,000) or PSER129 (diluted 1:50,000) antibodies diluted in blocking buffer overnight at 4°C. The following day membranes were washed 3 × 10 min in TBST and incubated with anti-rabbit (Invitrogen, 1:20,000) or anti-mouse (Cell Signaling, 1:6,000) HRP conjugates for 1 h at room temperature. Membranes were then washed 3 × 10 min in TBST. Membranes were imaged using ECL substrate and chemiluminescence imager (Biorad). 5 microliters of a broad range molecular weight standard (Biorad, product # 1610394) was used to determine approximate molecular weight of separated proteins.

### BAR analysis

Total normalized spectra values were exported from Scaffold. Heatmaps were generated using R with the heatmaply package. Differential expression analysis was performed and subsequent volcano plot generated using Graphpad Prism. Multiple paried t tests adjusted for multiple comparisons using a false-discover rate (FDR) was performed using a Two-stage step-up method of Benjamini, Krieger and Yekutieli with an FDR cutoff of 1% was performed. For this test, two conditions groups were compared, BAR-NEG vs. BAR-PSER129. BAR-PSER129 enriched proteins were compared across groups using venn analysis (Molbiotools.com, Multiple List Comparator). High-confidence functional and physical protein-protein interactions were plotted using STRING with MCL clustering (inflation parameter 1.4). Clusters were annotated with notable significant pathways enrichments.

### Quantification of olfactory bulb layer pathology

To quantify αsyn pathology in the OB, three representative bright field images of each OB layer (i.e., GL, mitral cell layer, outer plexiform layer, and glomerular layer) for each animal were taken using 20X objective. Images were then masked using the Threshold algorithm in Nikon elements. Using methyl green counterstain as a guide, R0I’s were drawn over each layer of the OB and the mean density of this area calculated. Density values were exported and organized in Excel (Microsoft) and then graphed using GraphPad. All Images were gathered and quantified by a reviewer blinded to experimental treatment information.

### Graphing and Statistical Analysis

Graphs were created using GraphPad Prism. For behavioral analyses and most neuroanatomical analyses described above, a 2-way repeated measures analysis of variance will be performed with group and time being the main variables. If there is a significant group by time interaction, we will perform post-hoc tests that control for multiple comparisons. If a non-normal distribution of data is determined, a Friedman’s one-way ANOVA will be employed for non-parametric data and if significant, would be followed by a Mann-Whitney U test for individual comparisons.

## Supporting information

Case Table

## Acknowledgements

Proteomics services were performed by the Northwestern Proteomics Core Facility, generously supported by NCI CCSG P30 CA060553 awarded to the Robert H Lurie Comprehensive Cancer Center, instrumentation award (S10OD025194) from NIH Office of Director, and the National Resource for Translational and Developmental Proteomics supported by P41 GM108569. RM lab was supported by France Parkinson and EraPerMed DEEPEN-iRBD project (ANR-22-PERM-0006). ES is supported by the Intramural Research Programs of the National Human Genome Research Institute and National Institutes of Health. Additionally, she also receives funding from the Aligning Science Across Parkinson’s (ASAP-000458) Michael J. Fox Foundation and a Collaborative Research Agreement between the NHGRI and Roche (Basel). BAK receives support from NINDS award #1R01NS128467 and Michael J. Fox Foundation. This work was supported in part by Aligning Science Across Parkinson’s [ASAP-024442] through the Michael J. Fox Foundation for Parkinson’s Research (MJFF) and NIH R21 NS109871 to JHK. GP received support from NINDS grant # K23-NS097625-06.

