## Supplemental Figures for "Neither alpha-synuclein-preformed fibrils derived from patients with *GBA1* mutations nor the host murine genotype significantly influence seeding efficacy in the mouse olfactory bulb"

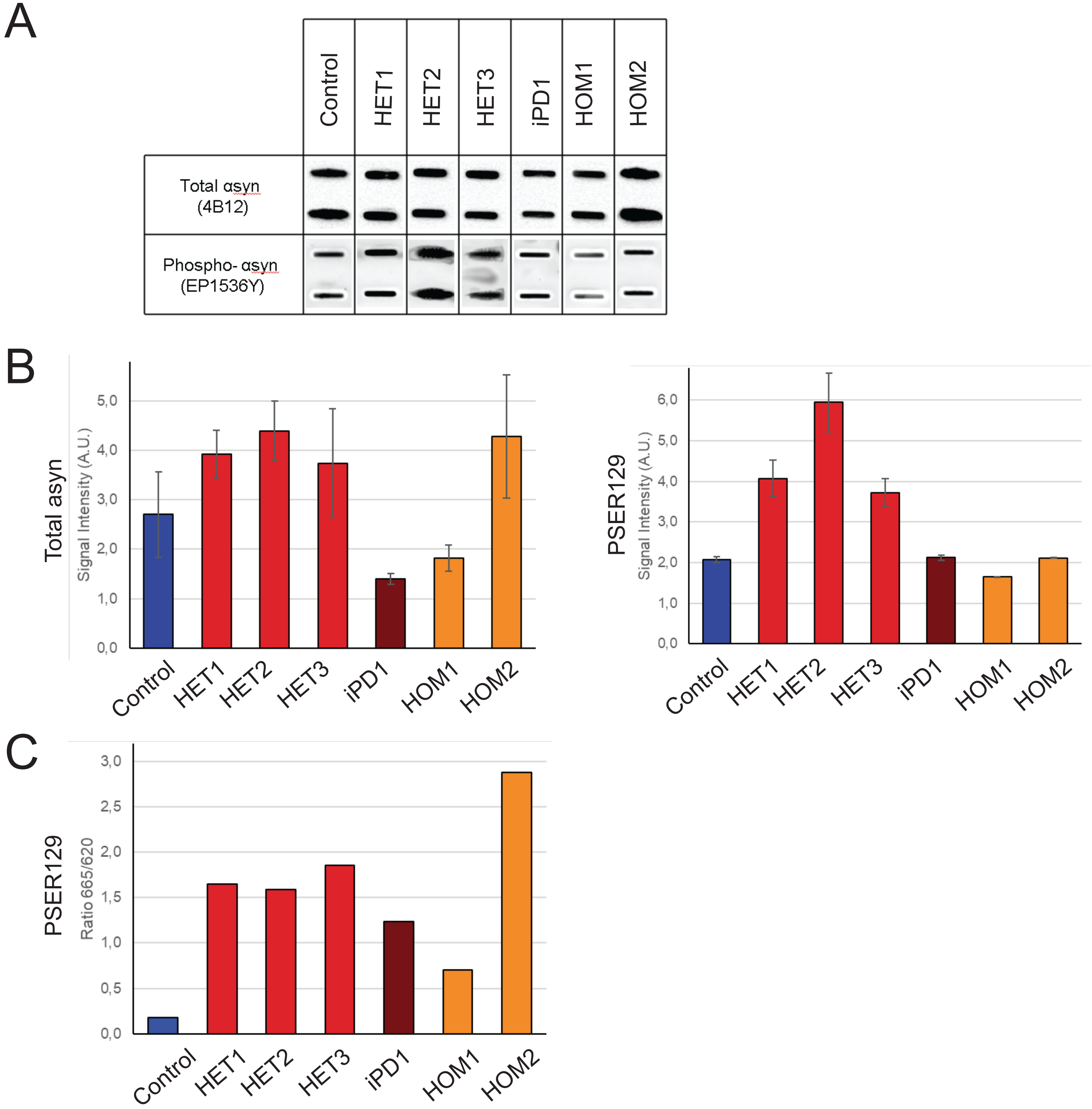


**Figure S1. Characterization of patient brains homogenates.**
(A) The amount of total and phosphorylated αsyn in the different brain homogenate was determined using a filter retardation assay. 50µL of brain homogenates (1% W:V) were filtered in duplicate on nitrocellulose membrane and probed with 4B12 (total αsyn) or EP1536Y (phosphorylated αsyn). (B) Quantification of total and phosphorylated αsyn in the different brain homogenate presented in panel A, bars represent ±SD. (C) The amount of pathogenic phosphorylated αsyn in the different brain homogenates (2,5% W:V) was quantified using the cisbio FRET assay following the manufacturer's recommendations.

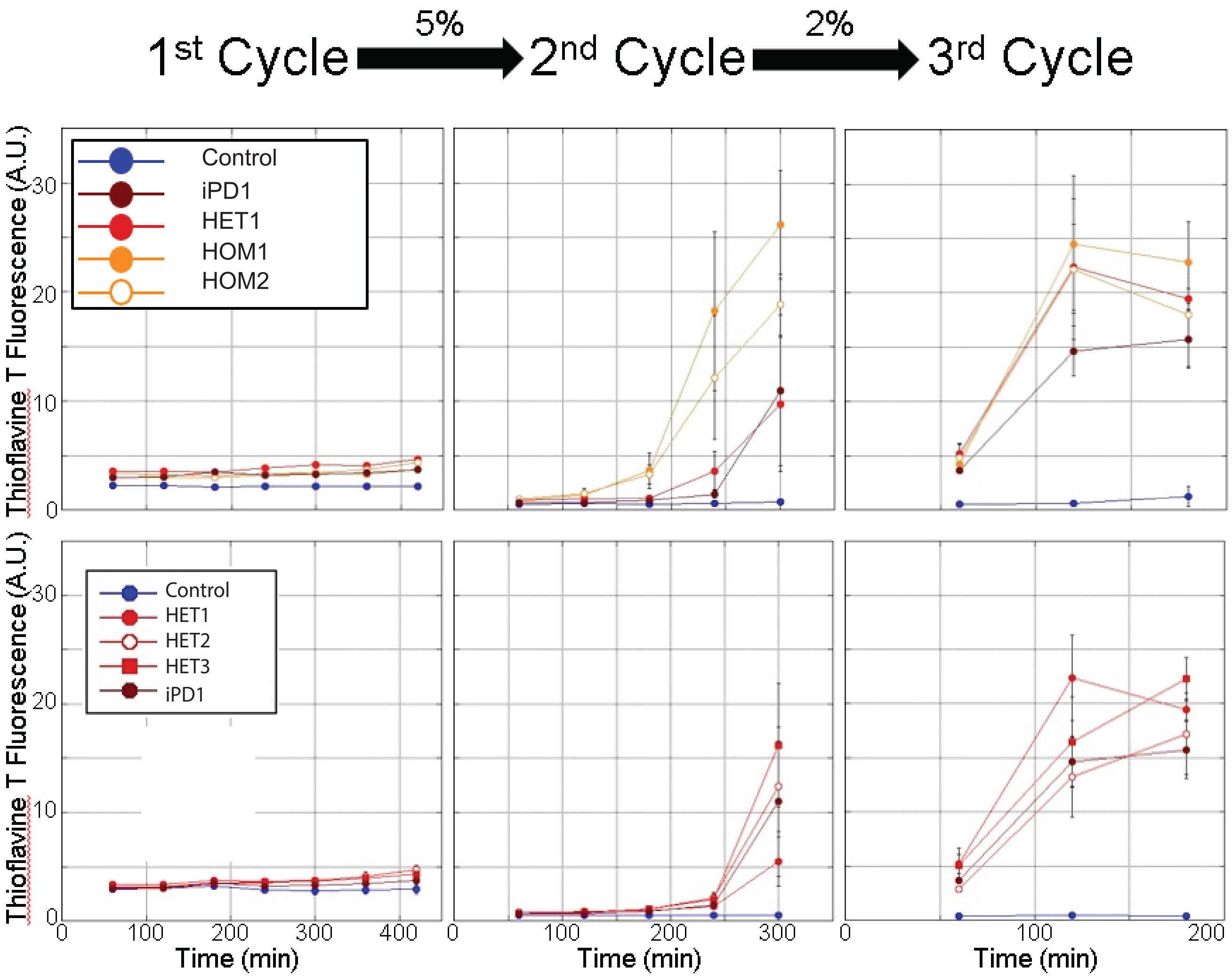


**Figure S2. Amplification of pathogenic αSYN from patients brain homogenates by PMCA.** PMCA was performed on human brain homogenates (2% (W:V) for the 1st cycle, the indicated amounts (V:V) for the next cycles, in PMCA buffer containing monomeric αsyn (100 μM). The amounts of brain homogenates and PMCA-amplified assemblies used in each amplification reaction were optimized through several trials to minimize the de novo aggregation of αsyn under our experimental conditions. The time at which an aliquot from one amplification reaction was withdrawn for a subsequent amplification reaction was also optimized to avoid the formation of de novo of αsyn fibrillar assemblies. The curves represent an average of four replicates ± SD

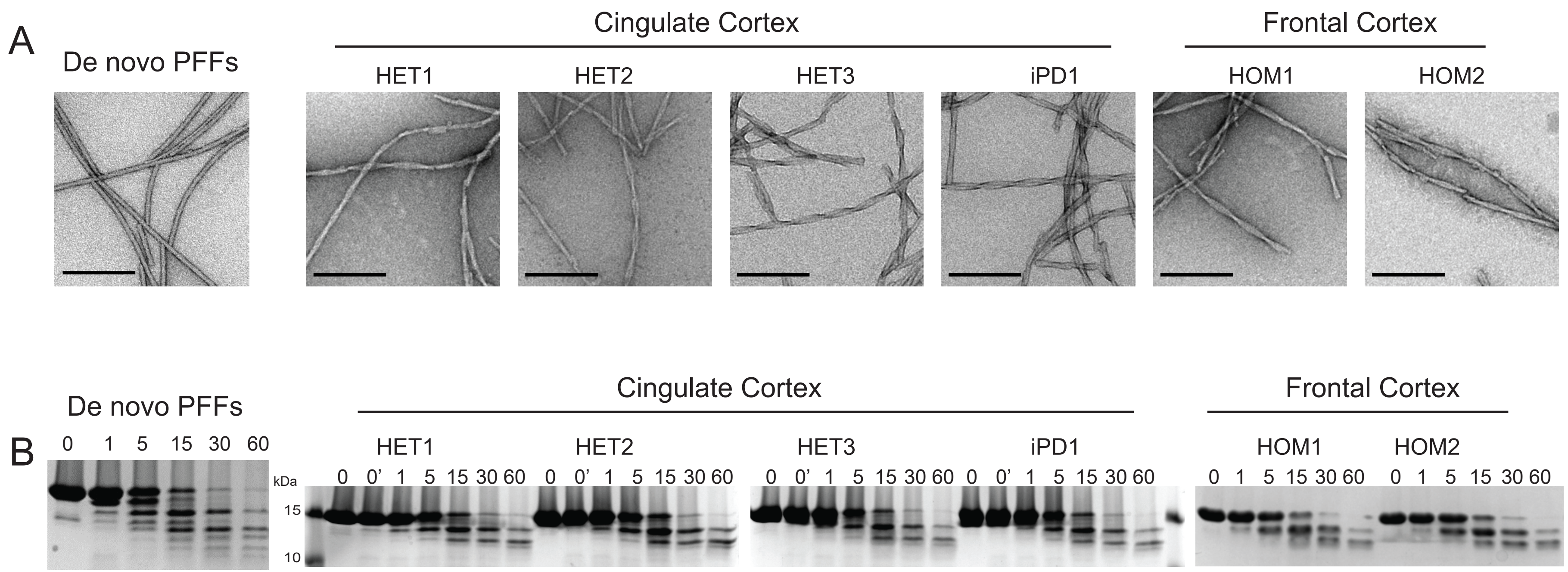


**Figure S3. In vitro characterization of patient-derived αsyn PFFs.**
(A) Electron micrographs of of patient-derived αsyn PFFs obtained after the 4th cycle of amplification by PMCA and de novo generated PFFs. Scale bar = 200 nm. (B) Limited proteolytic patterns of the different strains. Monomeric αsyn concentration is 100 μM. Proteinase K concentration is 3.8 μg/ml. Samples were withdrawn from the reaction before PK addition (lane most to the left), immediately after PK addition (second lane from left) and at time 1, 5, 15, 30 and 60 min from left to right in all panels. PAGE analysis was performed and the gels were stained with Coomassie blue. The position of the molecular weight markers 15 and 10 kDa is indicated on the left.

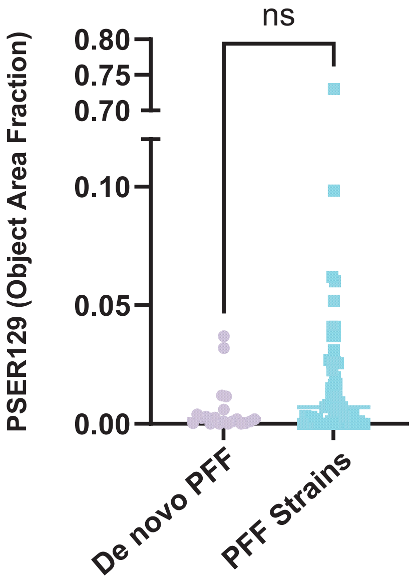


**Figure S4. PSER129 abundance in mice seeded with de novo PFFs or PFF strains.** ns= non-significant using Unpaired Two-tailed t test.

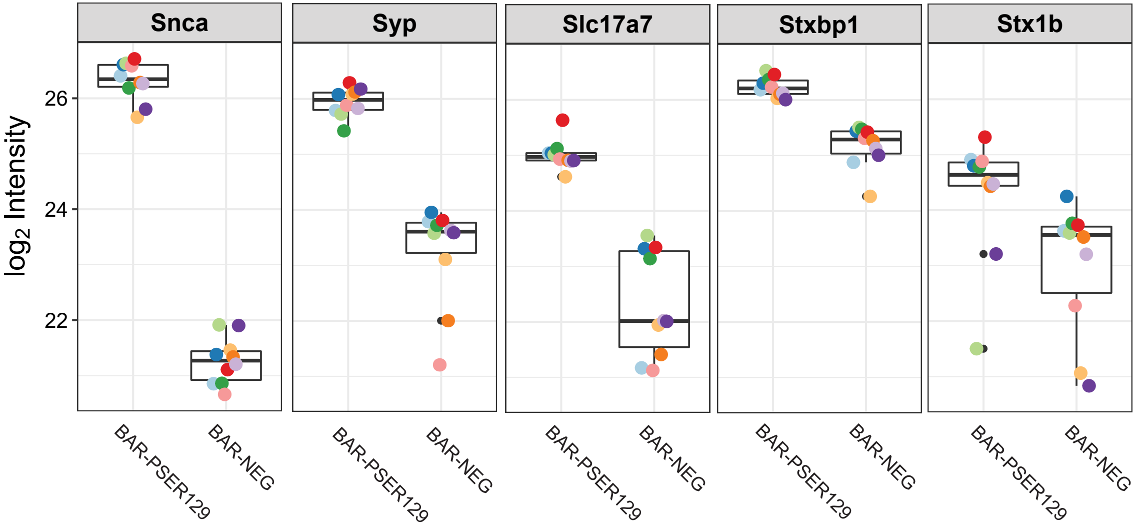


**Figure S5. BAR target enrichement.** .Raw LC-MS/MS data files were analyzed by Maxquant and LFQ-Analyst (Shah AD, Goode RJA, Huang C, Powell DR, Schittenhelm RB. LFQ-Analyst: An easy-to-use interactive web-platform to analyze and visualize proteomics data preprocessed with MaxQuant. DOI: 10.1021/acs.jproteome.9b00496). Two experimental groups were compared, BAR-Neg and BAR-PSER129. Boxplots of abundance show selected significant proteins including BAR target asyn (SNCA), and several known presynaptic vesicle/SNARE proteins thought to be closely associated with asyn. Results demonstrate high enrichment of asyn and the presynaptic compartment using BAR-PSER129. SYP = Synaptophysin, Slc17a7 = Vesicular glutamate transporter 1, Stxbp1 = Syntaxin binding protein 1, Stx1b = Syntaxin-1B1.

| Case id | Study Group | Age (y.) | Sex | GBA mutation | Gaucher pathology | Diagnosis | Duration (y.) | Braak Stage | Brain Region |
|---|---|---|---|---|---|---|---|---|---|
| HET1 | $GBA^{-/+}$ | 63 | F | H255Q | Unknown | Unknown | Unknown | 2 | Cingulate |
| HET2 | $GBA^{-/+}$ | 74 | M | E326K | Unknown | Unknown | Unknown | 3 | Cingulate |
| HET3 | $GBA^{-/+}$ | 85 | F | L444P | Unknown | PD/Dementia | Unknown | Unknown | Cingulate |
| iPD1 | $GBA^{+/+}$ | 93 | M | NA | NA | iPD | 12 | Unknown | Cingulate |
| HOM1 | $GBA^{-/-}$ | 61 | Female | N370S/c.84insG | Severe | DLB | 5 | Unknown | Frontal cortex |
| HOM2 | $GBA^{-/-}$ | 73 | Female | N370S/N370S | Minimal | DLB | Unknown | Unknown | Frontal cortex |

**Table S1. Case information.**

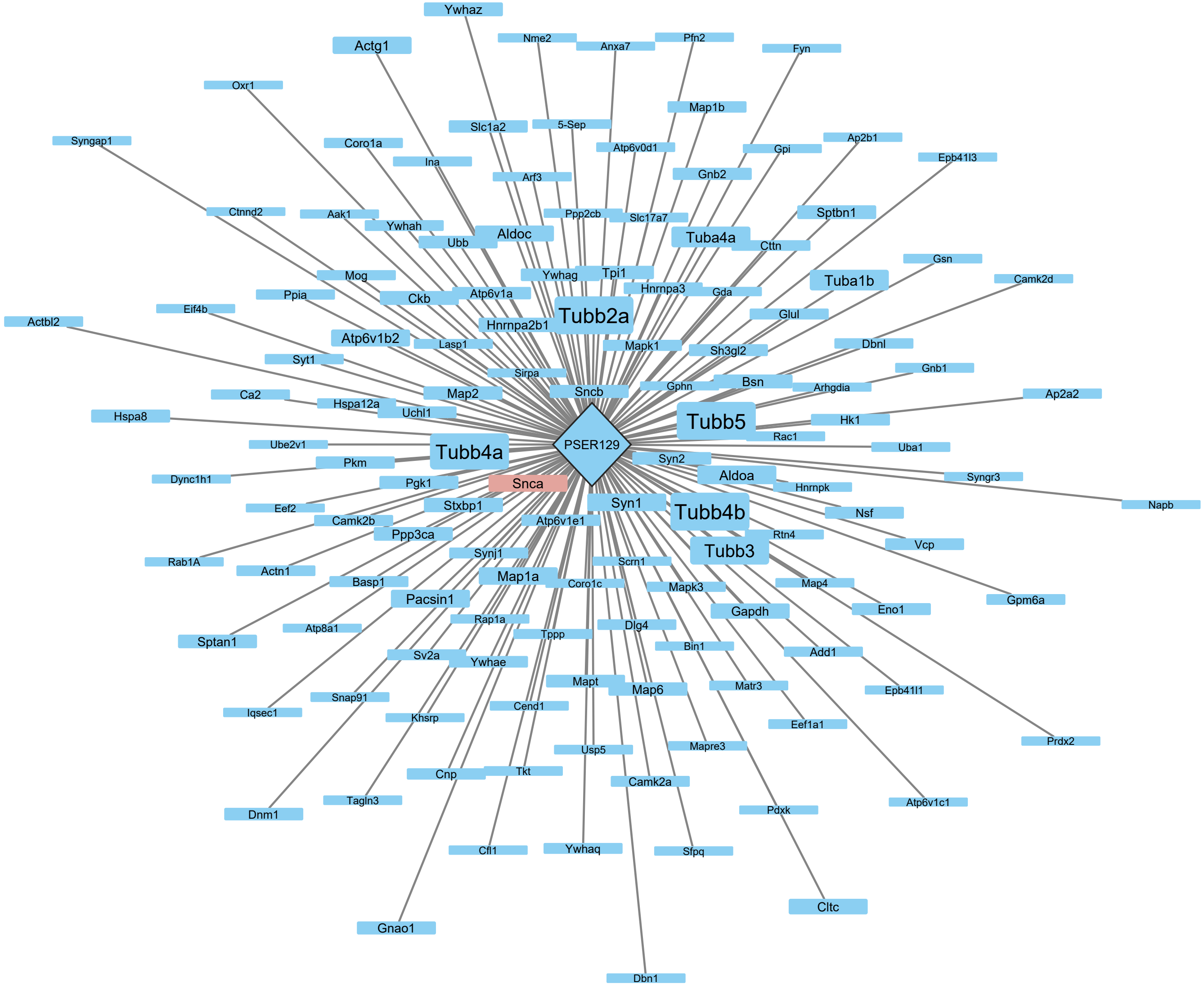


**Figure S6. Summary of BAR-PSER129 Enriched Proteins.** Enriched proteins included in Fig. 5 were plotted using cytoscape and perfusion force directed layout. Node and text size is directly proportional to difference for total spectra between BAR-PSER129 and BAR-NEG for each protein. The edge length, thus distance from center node “PSER129” is directly proportional to the corresponding q-value for each enriched protein. Alpha-synuclein’s node position is highlighted red.

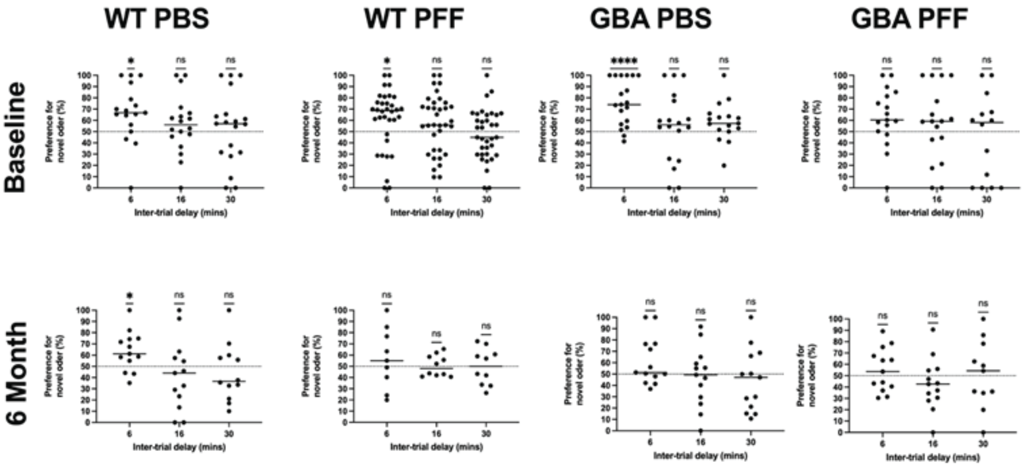


**Figure S7**: Odor retention test in GBA1$^{D409V/D409V}$ mice treated with different alpha-synuclein human pre-formed fibrils (HuPFFs). At baseline odor retention, the WT mice injected with PBS and HuPFFs and the GBA1$^{D409V/D409V}$ mice injected with PBS showed preference for the novel odor up until 6 mins. This suggest that they remembered the familiar odor for 6 mins, but did not remember it at 16 or 30 mins after initial exposure. The GBA1$^{D409V/D409V}$ mice injected with HuPFFs did not show preference for the novel odor at any time point, suggesting that they did not remember the familiar odor from the start.

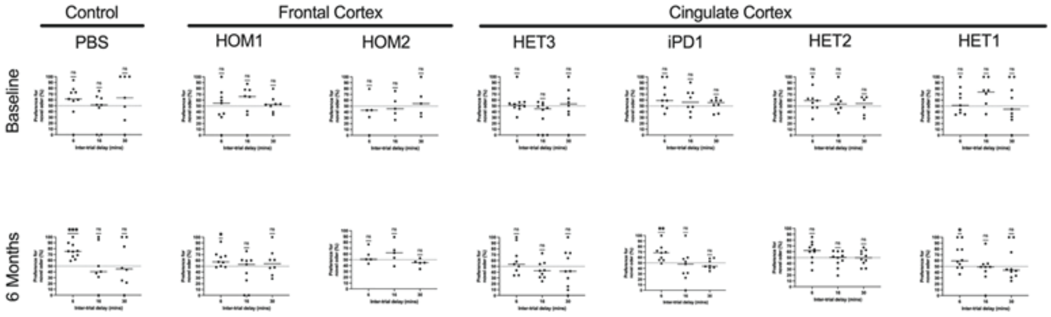


**Figure S8**: Odor retention test in wild type mice treated with different alpha-synuclein polymorphs (strains). The WT mice injected with PBS, GBA-HuPPFs (HOM1, HOM2, HET1, HET2, HET3), or iPD-HuPFFs (iPD1) did not show a preference for the novel odor during the baseline test. At 6 months, mice treated with B19 and GBA1 variant fibrils had a preference for the novel odor up until 6 mins after the initial exposure. This data suggests that, although there was some recognition at 6-month time points, short-term olfactory memory was impaired in the GBA-HuPPF and iPD-HuPFF injected mice.

A

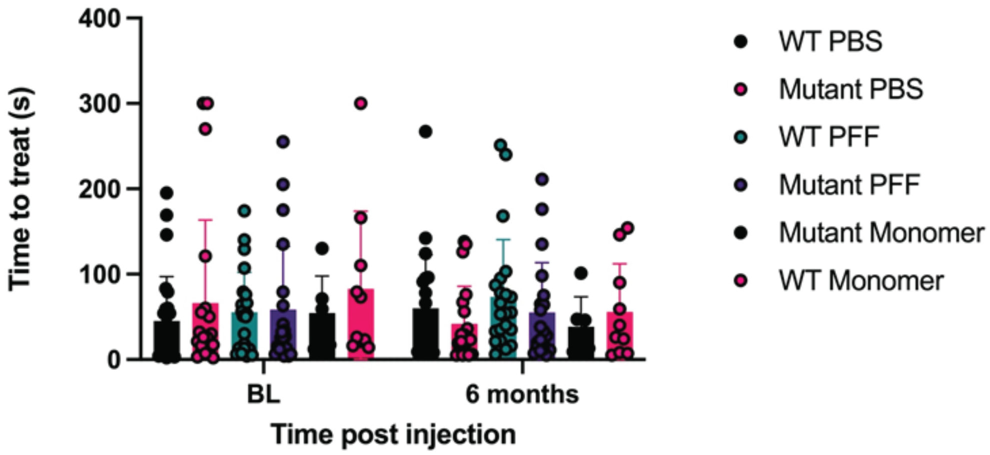


B

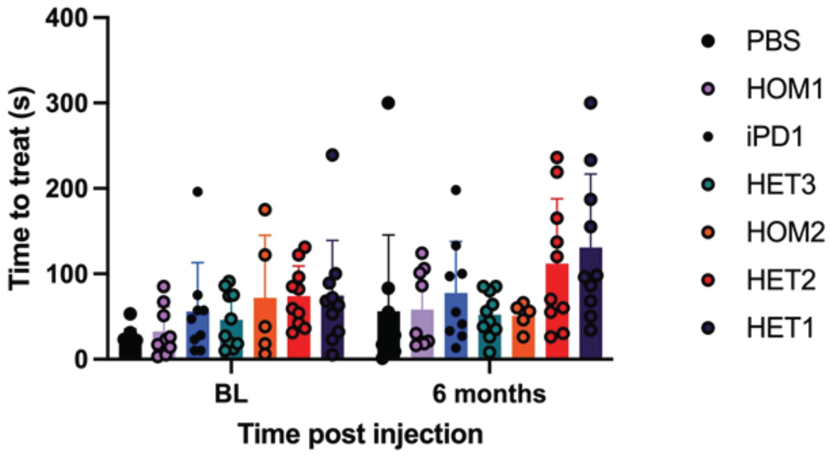


**Fig S9**: Digging odor test in GBA1$^{D409V/D409V}$ mice treated with different alpha-synuclein human pre-formed fibrils (hPFFs), (A) and wild type mice treated with different alpha-synuclein polymorphs (strains) (B). There was no change in olfactory function observed in the GBA1$^{D409V/D409V}$ mice over the course of 6 months post-injection. This suggests that genotype and WT α-syn fibril injection did not affect olfactory function. There was also no loss of olfactory function over the course of 6 months for the GBA-HuPPF and iPD-HuPFF injected mice, suggesting that α-syn from GBA mutation carriers does not impair olfactory function more than α-syn from iPD patients.

A

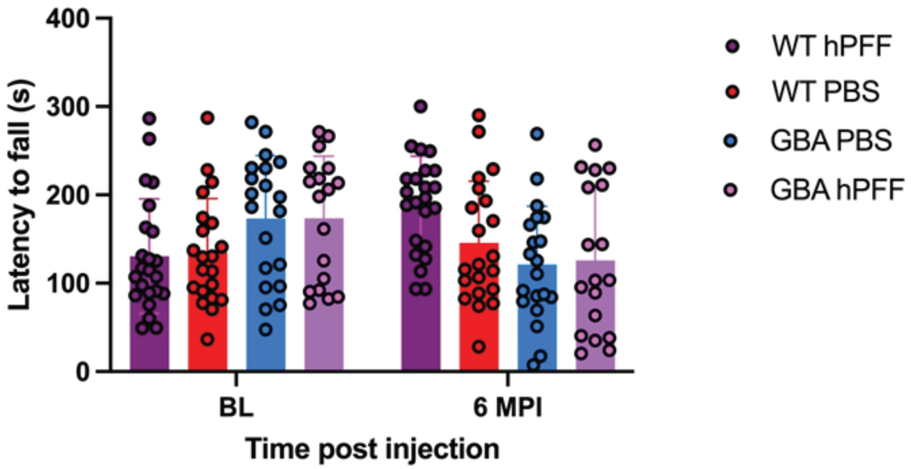


B

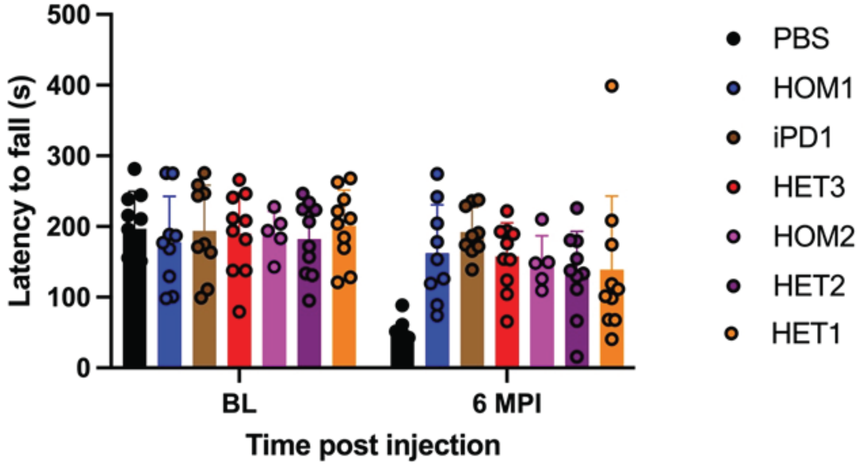


**Fig S10**: Rotarod test in GBA1$^{D409V/D409V}$ mice treated with different alpha-synuclein human pre-formed fibrils (hPFFs), (A) and wild type mice treated with different alpha-synuclein polymorphs (strains) (B). There was no argument for motor phenotype caused by HuPFFs or genotype based on the accelerating rotarod test. However, as there was no change in the GBA1$^{D409V/D409V}$ mice, it could be inferred that these mice failed to learn how to stay on the rotarod. This could suggest that there was cognitive impairment in these mice.

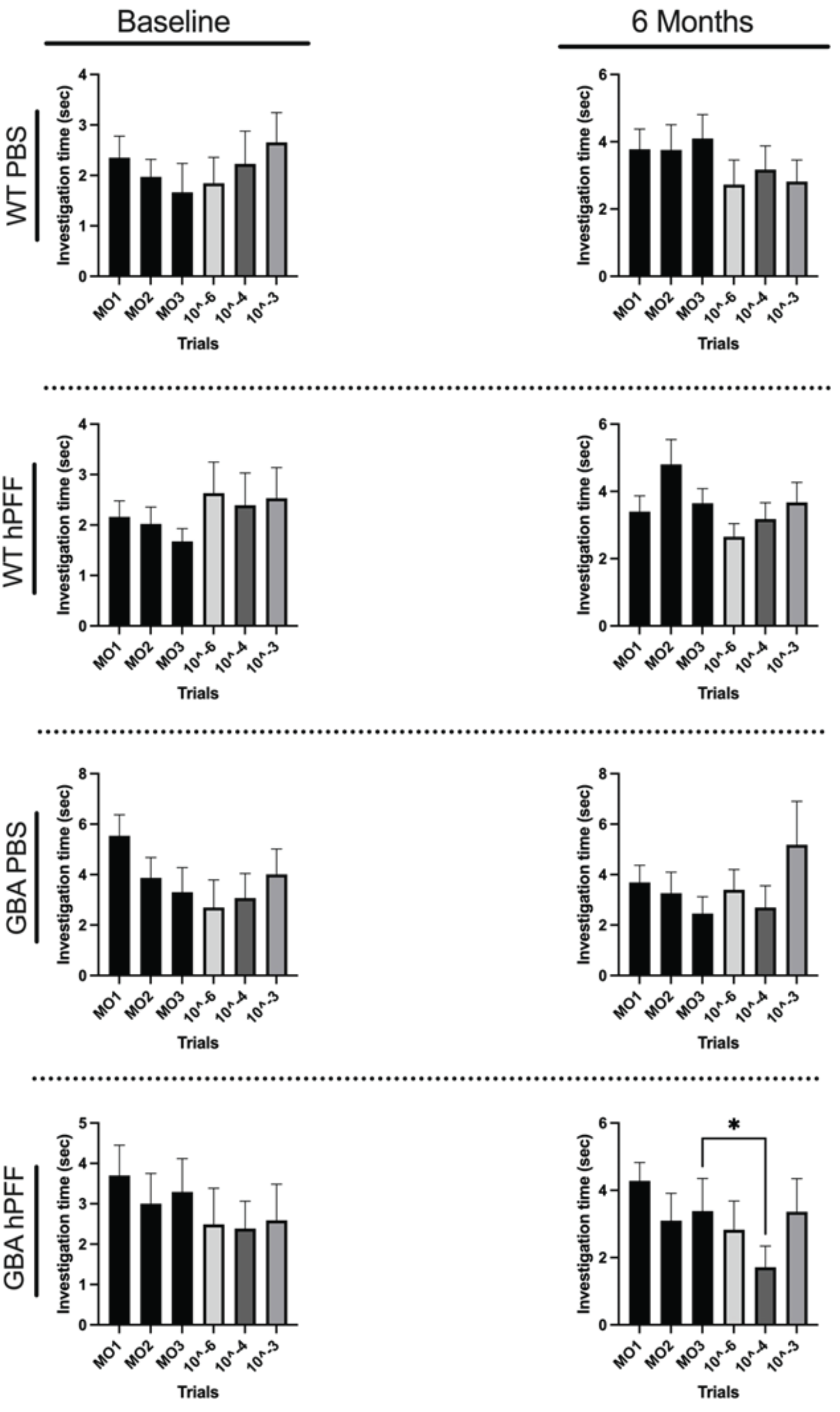


Fig S11: Odor threshold test in GBA1$^{D409V/D409V}$ mice treated with different alpha-synuclein human pre-formed fibrils (hPFFs). At baseline odor threshold, the WT and GBA1$^{D409V/D409V}$ mice that were injected with either PBS or WT-HuPFFs, did not show any ability to detect propionic acid at the three different concentrations (1:10$^{6}$, 1:10$^{4}$, and 1:10$^{3}$). This continued up until 6 months.

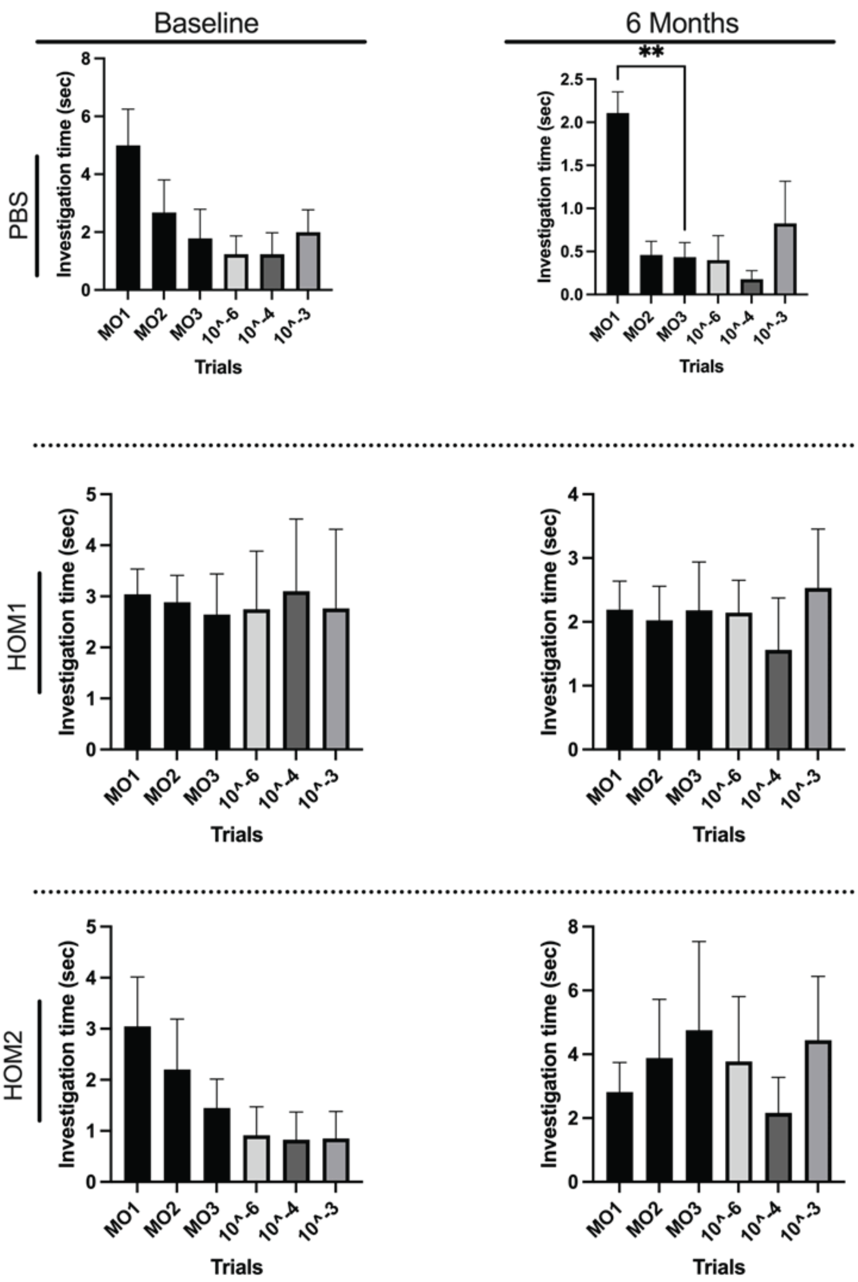


**Fig S12**: Odor threshold test in wild type mice treated with different alpha-synuclein polymorphs (strains) extracted from the frontal cortex. HOM1 and HOM2 are GBA-HuPPFs. The mice injected with PBS or GBA-HuPPFs did not show any ability to detect propionic acid at the three different concentrations (1:$10^6$, 1:$10^4$, and 1:$10^3$) from baseline to 6 months.

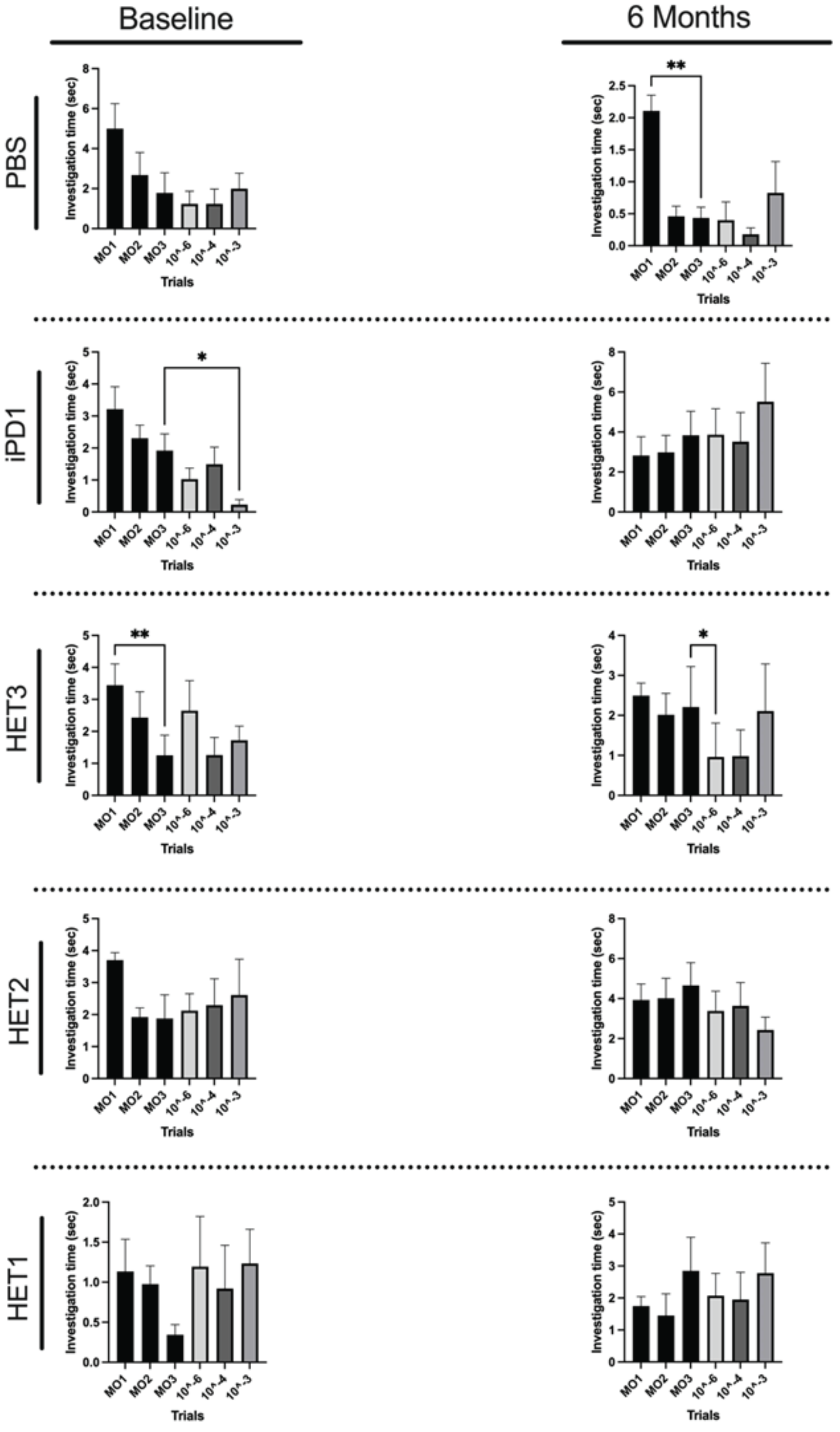


**Figure S13**: Odor threshold test in wild type mice treated with different alpha-synuclein polymorphs (strains) extracted from the cingulate cortex. The mice injected with PBS, GBA-HuPPFs (HET1, HET2, HET3) or iPD-HuPFFs (iPD1) also did not show any ability to detect propionic acid at the three different concentrations (1:10$^{6}$, 1:10$^{4}$, and 1:10$^{3}$) from baseline to 6 months.
