## Supplemental methods for "Neither alpha-synuclein-preformed fibrils derived from patients with *GBA1* mutations nor the host murine genotype significantly influence seeding efficacy in the mouse olfactory bulb"

**Odor retention test**

The odor retention test assesses short-term olfactory memory using pairs of odorants and is based on methods adapted from previous work [1, 2]. Before the first test, mice were pre-habituated to the setup for 5 min with empty cartridges in a clean mouse cage without bedding. During the test (performed in a blinded manner), the mice were exposed to two separate cartridges at a time that each contained a paper swab with an odorant infused. During the first trial (Acquisition), mice were exposed to the same unfamiliar odorant in the two cartridges. After 6, 16, or 30 min, mice were put through a second trial (Recall) that contained a cartridge containing the same odorant as the first trial, and a second cartridge with a novel odorant. The preferred odor was recorded and the preference was calculated for the recall trial. Three pairs of odorants were used (familiar and novel odor) at supraliminal concentrations. The chosen odorants were known for having equal hedonicity among each pair [1,3] (-) limonene (diluted 1:10) and (+) carvone (diluted 1:10); Amylacetate (1:100) and Anisol (1:100); and propyl acetate (1:10) and Pentanal (1:10). These odorants were diluted in mineral oil. The use of each odorant from the pair as familiar or novel odor was randomized. Data was expressed as a preference for the novel odor. The mean preference for novel odor for each group during the recall trial was analyzed by one-sample Student’s t test compared with the chance level of 50% (Prism 6.0; GraphPad). The odorants used were purchased from Sigma-Aldrich, TCI America, or provided by A. Didier (Lyon Neuroscience Research Center, Lyon, France).

**Odor Threshold**

The animals were placed a clean mouse cage without bedding. The test was done individually and each mouse was habituated to the setting during a pre-habituation trial with a cartridge that contained a paper swab soaked in mineral oil (MO). Each mouse was then exposed to a paper swap soaked in mineral oil for three 50s trials with a 5 min inter-trial interval (habituation phase, MO). The animals were then presented with a paper swab soaked in an odorant (propionic acid) at increasing concentrations (1:10^6^, 1:10^4^, and 1:10^3^, detection phase). in mineral oil. During each trial, the investigation time, defined as the duration of active sniffing with the nose placed less than 1 cm away from the cartridge. Mice that did not investigate the mineral oil during the first habituation trail were excluded. The mean investigation time per trial was calculated and anylyzed by one-way ANOVA with repeated measures across trials for each group and time point. This was done for the habituation phase first which is critical for being able to interpret the results from the following detection phase, and then for the detection phase. This was followed by a Sidak post-hoc test (more conservative than Fisher LSD post-hoc test) to compare MO1 to MO2, MO3, and MO3 to odorant concentrations 1:10^6^, 1:10^4^, and 1:10^3^ using Prism 6.0 (GraphPad Software).

**Rotarod**

The rotarod test was used to assess motor learning, coordination, and balance in the mice (MED-Associates). Each mouse was given a training session (four 5- min trials, 5 min apart) to acclimate them to the rotarod apparatus. During the test period (1 hr later), each mouse was placed on the rotarod with increasing speed, from 4 rpm to 40 rpm in 300 sec. The latency to fall off the rotarod with in this time period was recorded. Each mouse received two consecutive trials and the mean latency to fall was used in the analysis.

**Digging Test**

Prior to testing animals were fasted overnight. Fasting did not to exceed 24h. A clean mouse cage (15 cm x 3.25 cm x 3.13 cm) was filled with 3 cm of bedding and an appetitive stimulus of sweetened cereal (Cinnamon Toast Crunch) buried 0.5 cm below the bedding and along the perimeter of the cage. The animal were monitored for 5 mins or until the animal found the food treat (latency to find treat), at which time the session was completed. Once the session was completed, the animals were returned to their home cage and food returned. If the mouse found the treat, it was allowed to eat it. If the treat was not found within 5 mins the mouse placed back in its home cage and the treat was removed. The bedding was changed between mice.
